# RNA-guided *As*Cas12a- and *Sp*Cas9-catalyzed knockout and homology directed repair of the *omega-1* locus of the human blood fluke, *Schistosoma mansoni*

**DOI:** 10.1101/2021.11.15.468743

**Authors:** Wannaporn Ittiprasert, Chawalit Chatupheeraphat, Victoria H. Mann, Wenhui Li, André Miller, Taiwo Ogunbayo, Kenny Tran, Yousef Alrefaei, Margaret Mentink-Kane, Paul J. Brindley

## Abstract

We compared the efficiency of gene knockout (KO) and precision of insertion (knock-in, KI) of the RNA-guided *As*Cas12a nuclease of *Acidaminococcus* sp. with that of *Sp*Cas9 from *Streptococcus pyogenes*, aiming to enhance the functional genomics toolkit for *Schistosoma mansoni*. Programmed DNA cleavages catalyzed by Cas12a and Cas9 result in staggered and blunt ended strand breaks, respectively. TTTV, the optimal protospacer adjacent motif for *As*Cas12a would occur frequently within the AT-rich genome of this platyhelminth. We deployed optimized conditions for the ratio of guide RNAs to the nuclease, donor templates, and electroporation parameters, to target a key enzyme termed omega-1 that is secreted by the schistosome egg. *As*Cas12a was more efficient than *Sp*Cas9 for gene knockout of *omega-1* as determined by tracking of indels by decomposition (*P* < 0.001). Resulting from CRISPREsso2 analysis, most mutations were deletions; *Sp*Cas9 induced short deletions of 3 nt in length whereas *As*Cas12a induced deletions of 2 to 26 nt. Knockout efficiency of both nucleases markedly increased in the presence of short, single stranded oligodeoxynucleotide (ssODN) donor templates. With *As*Cas12a, ssODNs representative of both the non-CRISPR target (NT) and target (T) strands of the targeted gene were tested, resulting in KO efficiencies of 15.67, 28.71 and 21.43% in the *Sp*Cas9 plus donor ssODN, *As*Cas12a plus NT-ssODN, and *As*Cas12a plus T-ssODN groups, respectively. *Trans* cleavage activity against the ssODNs by activated *As*Cas12a was not apparent *in vitro*. Programmed *Sp*Cas9 editing led to more precise transgene insertion than *As*Cas12a, with KI efficiencies of 17.07% for the KI_SpCas9 group, 14.58% for KI_*As*Cas12a-NT-ssODN and 12.37% for KI_*As*Cas12a-T-ssODN. Although *As*Cas12a induced fewer mutations per genome than *Sp*Cas9, the phenotypic impact on transcription and expression of omega-1 was similar for both nucleases. These findings revealed that *As*Cas12a and *Sp*Cas9 both provide tractable routes for RNA-guided programmed mutation of the genome of the schistosome egg.

## Introduction

Interest is increasing in functional genomics to investigate helminth parasites, specifically genome editing to investigate pathophysiology, carcinogenesis, and biology. Recent progress showcased the deployment of CRISPR/Cas programmed genome editing in parasitic flatworms, in particular with the human blood fluke *Schistosoma mansoni* (Ittiprasert *et al*, 2019) and the carcinogenic liver fluke, *Opisthorchis viverrini* (Arunsan *et al*, 2019) as exemplars. However, given that these helminth parasites are non-model organisms, that their genome sequences are often not available or well curated, and that they are frequently refractory to maintenance in the laboratory, approaches to advance functional genomics using CRISPR/Cas are not yet established or available for wide use in the field of tropical medical and neglected tropical diseases. Nonetheless, our capacity to decipher the molecular pathogenesis of these diseases will be aided by advances in gene editing methodologies (Hoffmann *et al*, 2014).

RNA-guided nucleases evolved in archaea and bacteria as adaptive immunity mechanism to destroy phage viruses and other nucleic acid invaders (Jinek *et al*, 2012; Koonin *et al*, 2017). The catalogue of these CRISPR/Cas nucleases is expanding. The *Sp*Cas9 of *Streptococcus pyogenes*, a Class 2, type II Cas enzyme in the phylogeny of Koonin and coworkers (Koonin *et al*., 2017), dominates diverse gene-editing applications in biomedicine and beyond (Charpentier, 2015; Huang *et al*, 2018; Mustafa & Makhawi, 2020), given its programmable, precise double-stranded DNA (dsDNA) cleavage, accomplished by two catalytic domains, RuvC and HNH, of the nuclease at the target site complementary to the guide RNA (Nishimasu *et al*, 2014). A second Class 2 family, the type V nucleases as represented by *As*Cas12a (formerly known as Cpf1) uses a single RuvC catalytic domain for guide RNA-directed dsDNA cleavage (Jeon *et al*, 2018; Zetsche *et al*, 2015). Unlike *Sp*Cas9, *As*Cas12a orthologues recognize a T nucleotide–rich protospacer-adjacent motif (PAM), catalyze the maturation of their own guide CRISPR RNA (crRNA) and a dsDNA break, distal to the PAM, bearing staggered 5’- and 3’-ends (Jeon *et al*., 2018). These Cas12a features are attractive for genome editing (Malzahn *et al*, 2019). For their maturation, the crRNAs of type II CRISPR-Cas systems require processing by a cognate trans-activating CRISPR RNA (Charpentier *et al*, 2015). Moreover, the *As*Cas12a nuclease is smaller in size than *Sp*Cas9 and uses a shorter CRISPR RNA (crRNA) for activity (Liu *et al*, 2017). Notably, target site activated *As*Cas12a exhibits non-specific *trans* activity against bystander nucleic acids (Chen *et al*, 2018; Smith *et al*, 2020), an attribute that has enabled its application in a burgeoning range of biosensors (Kellner *et al*, 2019; Li *et al*, 2018a).

Delivery of CRISPR reagents into the cells of the multicellular parasite *S. mansoni* in the form of ribonucleoprotein (RNP) complexes may enable immediate and permanent programmed gene editing and obviate concerns with CRISPR reagent longevity or integration in the host cell genome. Programmed CRISPR/Cas editing using RNP-based delivery can precede analysis of mutated target regions that rely on cloning and vector construction. Previously, we deployed both RNP and lentiviral-delivered CRISPR/*Sp*Cas9 approaches to knock-out (KO) and knock-in (KI) into a blunt end at *Sp*Cas9 cleavage site at the multiple copies of the omega-1 (*ω1*) gene in parasite eggs. To advance functional genomics for schistosomes and aiming to improve the mutagenesis and transgene knock-in of CRISPR efficiency in schistosomes, which exhibit AT-rich genomes, we investigated the nuclease activity of *As*Cas12a in comparison with *Sp*Cas9. These CRISPR systems differ in key ways: first, *Sp*Cas9 recognizes NGG as the protospacer adjacent motif (PAM) whereas *As*Cas12a recognizes the T-rich, TTTV, and second, catalysis by *Sp*Cas9 results in blunt double stranded DNA at three nucleotides 5’ to the PAM whereas *As*Cas12a results in a staggered cleavage at 23 nt (target strand; T) and 18 nt (non-target strand; NT) 3’ to the PAM (Banakar *et al*, 2020). Comparisons of these two RNA-programmed nucleases in the context of the *ω1* gene of *S. mansoni* may inform decisions for functional genomics manipulations aiming to enhance efficiency of homology directed repair (HDR) targeting genes expressed by the developing schistosome egg, and indeed other developmental stages (Ittiprasert *et al*., 2019; Sankaranarayanan *et al*, 2021; Takaki *et al*, 2021a; Takaki *et al*, 2021b).

## Results

### The omega-1 multicopy gene, guide RNAs, and single stranded DNA donor templates

The gene editing efficiency of *Sp*Cas9 and *As*Cas12a was compared in assays targeting the *ω1* gene which encoded a T2 ribonuclease that is secreted by the mature egg of *S. mansoni* into the surrounding host tissues and circumoval granuloma (Schwartz & Fallon, 2018). Specifically, the four copies of *ω1*, well annotated in WormBase Parasite (Bolt *et al*, 2018) (update of 16 September 2021) as hepatotoxic ribonuclease – Smp_334170.1, Smp_334170.2, Smp_334240, and Smp_333930, were targeted for design of guide RNAs (gRNA) and donor repair templates. These four copies reside on chromosome 1 of *S. mansoni*, with reverse strand positions at 3,980,364 to 3,984675, 3,992,964 to 3,995,246 and 3,908,953 to 3,911,250 (PRJEA36577), respectively. These copies share >99% nucleotide sequence identity, are ~2.3 kb in length, are highly AT rich (65%), and encode an enzyme of 127 (Smp_334170.2, Smp_334240 and Smp_333930) or 115 (Smp_334170.1) amino acid residues in length. Smp_333220, also located on chromosome 1 at 3,925,420-3,927,716 but lacking annotation on the *S. mansoni* annotated genome version 7, Year 2021, was also included. Smp_333220 shares >99% chromosomal and mRNA sequence identity with the three *ω1* copies (Fig S1B). Guide RNAs for *Sp*Cas9 and *As*Cas12a designed with the assistance of CHOPCHOP webtool both target the same region of exon 1 of *ω1*, disrupting its catalytic active site histidine (Luhtala & Parker, 2010), with the programmed cleavage sites for Cas in close proximity, separated by three nucleotides (Fig 1A and B).

**Figure 1.**
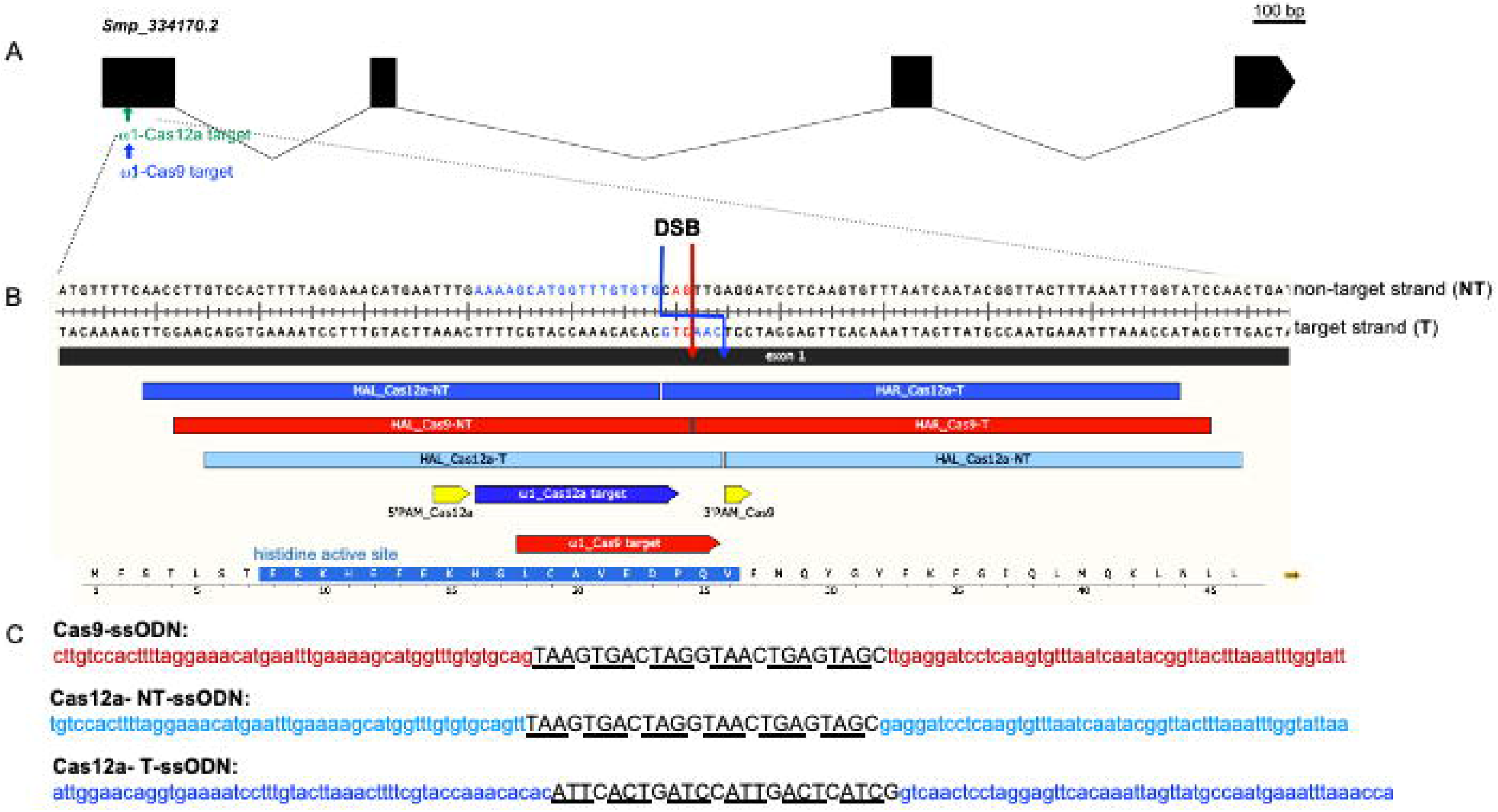
Schematic of the *Schistosoma mansoni* hepatotoxic ribonuclease omega 1 (ω1) gene structure, target sites for the *Sp*Cas9 and *As*Cas12a, ssODN donors and partial amino acid sequence with histidine active site. A Diagram of the omega-1 (*ω1*) gene (Gene ID; Smp_334170.2 of *S. mansoni* annotated genome version 7_2021) and CRISPR/Cas target sites on coding sequence of the first exon coding for histidine active site, FRKHEFEKHGLCAVEDPQV. The relative positions of crRNA for CRISPR-*Sp*Cas9 (red arrow) and -*As*Cas12a (blue arrow) are indicated along with their 3’PAM (yellow arrow) and 5’PAM (yellow arrow), respectively. B Programmed cleavage site for *Sp*Cas9/sgRNA ~3nt upstream of PAM 3’TGG (red arrow); and cleavage site for *As*Cas12a/crRNA at 18-24 nt downstream of a prospective 5-’TTTV PAM (blue arrow). The programmed cleavage sites for both enzymes, *Sp*Cas9 and *As*Cas12a, were three to six nt distant from each other. Fifty residues upstream and downstream of the double strand break (DSB) were included as the homology arms (HA) of the six stop codon cassette transgene (payload/cargo). Because cleavage by *Sp*Cas9 liberates blunt ended DNA, only a non-target strand (NT) repair template was provided for HDR (red bar). Sticky ended DNA results from cleavage by *As*Cas12a and hence both non target and target strand (T) repair templates for HDR were provided (light and dark blue bars). C The DNA donor sequences of CRISPR/*Sp*Cas9 and CRISPR/*As*Cas12a with non-target (NT) or target (T) sequence HA where the *As*Cas12a-NT-ssODN donor repairs the anti-sense DNA and *As*Cas12a-T-ssODN donor repairs the sense stand.

We designed three single stand oligodeoxynucleotide (ssODN) donors for homology directed repair (HDR) of *ω1* following programmed mutation. Given the differing PAM requirements of *Sp*Cas9 and *As*Cas12a as well as the divergent formats of the DSB, blunted ended by *Sp*Cas9 and sticky ended by *As*Cas12a, the designs differed for the ssODNs for use with each nuclease. Based on earlier findings (Ittiprasert *et al*., 2019), the design of the ssODN donors included 3’ and 5’ homology arms (HA) of each of 50 nt in length, complementary to the target strand for *Sp*Cas9 and for *As*Cas12a, for either the target strand, termed *Sp*Cas9-T-ssODN, or for the non-target strand donor, termed *As*Cas12a-NT-ssODN. Note that sequences identical to the PAM for *As*Cas12a and ~18 nt of the protospacer region were retained within the 5’-HA of *As*Cas12a-T-ssODN (Fig 1B and C).

### Absence of *trans* activity of activated *As*Cas12a against the ssODN donors

Previous reports dealing with *Lachnospiraceae bacterium* ND2006 and *Francisella novicida As*Cas12a describe indiscriminate *trans*-activity unleashed on ssDNA after gRNA activation and target binding of *As*Cas12a (Chen *et al*., 2018; Gier *et al*, 2020). Given that our goal here was to compare *Sp*Cas9 and *As*Cas12a in schistosomes, and given that we used an orthologous *As*Cas12a from *Acidaminococcus* sp. BV3L6 (IDT), in both KO and KI assays, we sought to confirm whether the omega-1 target gene activated *As*Cas12a might exhibit also *trans*-activity against the donor single stranded templates that we planned to use in KI assays, *As*Cas12a-T-ssODN and *As*Cas12a-NT-ssODN, in addition to RNA-guided double stranded DNA cleavage activity.

Accordingly, we also carried out *in vitro As*Cas12a nuclease activity assays using RNPs targeting a 426 bp amplicon of the exon 1 of *ω1* (Fig A, 2B and 2C). These assays confirmed that the crRNA was highly active and following incubation at 37°C for 30 min had cleaved the double stranded amplicon at the programmed cleavage site into two fragments of 265 and 161 bp, as expected (Fig 2B; Agilent Bioanalyzer electropherogram). In contrast, the activated *As*Cas12a-RNP complex failed to digest either of the single stranded templates, ssODNs *As*Cas12a-T-ssODN or *As*Cas12a-NT-ssODN after 120 min at 37°C (Fig 2C). As seen in the electropherogram, the ssODNs remained intact at 124 nt. Single stranded DNA standards ranging in size from <40 to 200 nt were included in the assay (Fig 2C). In summary, the findings suggested that *omega-1* target gene activated *As*Cas12a would not exhibit indiscriminate *trans*-nuclease activity against the ssODNs within schistosome cells, although an *in vitro* surrogate assay cannot faithfully mimic conditions within a viable schistosome egg.

**Figure 2.**
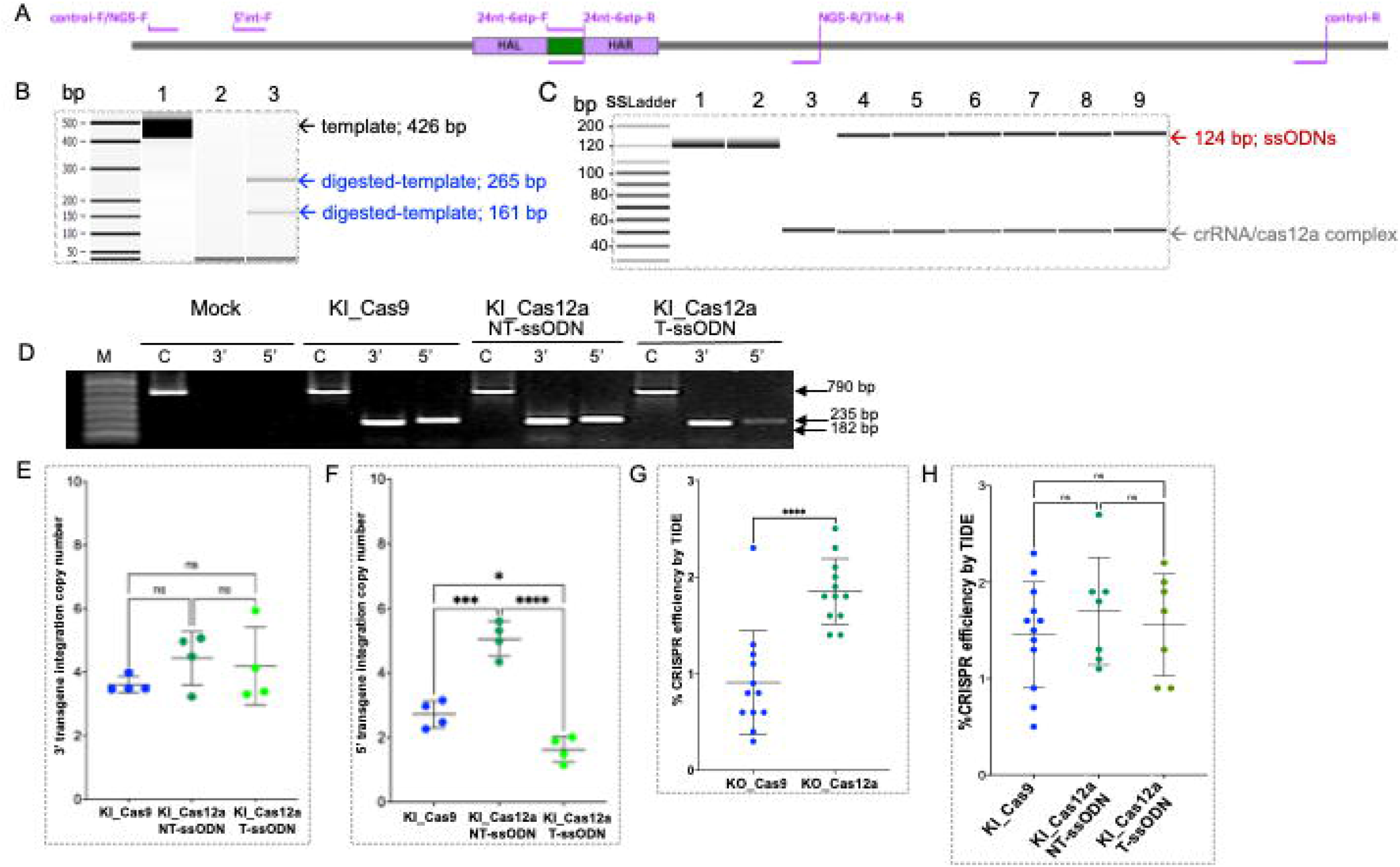
Knock-in by homology directed repair. A The location of primers used for 5’- and 3’-transgene (purple lines) knock-in PCR (5’int-F with 24nt-6stp-R and 24nt-6stp-F with 3’int-R) and internal positive control (control-F and -R). The primers for next generation sequencing indicate as NGS-F and NGS-R. All primers are located outside of the HA (purple boxes) sequences to avoid also positive amplification from DNA donors. The 24 nt-6stp-F and -R primers are highly specific for the stop codon transgene, and should not anneal to the wild type schistosome gene. B The *As*Cas12a target recognition activates specific 426 bp dsDNA amplicon template (lane 1) cleavage into 265bp and 161 bp (lane 3) products. The crRNA/*As*Cas12a RNP complex of ~40 nt is seen in lanes 2 and 3. C No specific ssDNA cleavage was seen on the Cas12a-NT-ssODN or Cas12a-T-ssODN donors at 124 nt as in lanes 1 and 2, respectively. The crRNA/*As*Cas12a RNP (~40 nt in lanes 3-9) containing activated *As*Cas12a did not show indiscriminate ssDNA digestion at 30 min, 60 and 120 min in lanes 4-5, 6-7 and 8-9, respectively. Lanes 4, 6 and 8 show the uncut 124 nt of Cas12a-NT-ssODN (top band) with Cas12a-crRNA (lower band), and lanes 5, 7 and 9 shows 124 nt of Cas12a-T-ssODN (top band) with Cas12a-crRNA (lower band). D The PCR detection of the integration sites in the transgenic eggs and control. The amplified control product (lanes C) for the wildtype sequence was 790 bp in length while those for the 5’(lanes 5’) and 3’ (lanes 3’) flanking regions of the transgenic sequence were 235 bp and 182 bp due to KI of Cas-9 ssODN, Cas12a_NT-ssODN and Cas12a_T-ssODN donor templates, respectively. Lane M is a 1 kb Plus DNA ladder; mock (only buffer electroporation) served as the negative control for transgene integration. E, F The estimation of transgene copy number in egg DNA were calculated by qPCR comparing with known quantity of the 24 nt transgene in 235 bp (5’ integration PCR fragment) ligated-pCR4-TOPO plasmid DNA (size 4,191 bp) standard curve; y = −4.0964ln(10) + 44.821, R^2^ = 0.9923 (data not shown). There was the confirmation of 3’ integration of 3.61±0.25, 4.44±0.84 and 4.19±1.22 of transgene copy from KI_Cas9, KI_Cas12a_NT-ssODN and KI_Cas12a_T-ssODN, respectively (one-way ANOVA, multiple comparisons, 95% CI of diff, *P*-value at 0.4057, 0.6264 and 0.9155, not significant). While asymmetric HDR was revealed by 5’ transgene integration of 2.72±0.42, 5.06±0.54 and 1.64±0.39 copy number from KI_Cas9, KI_Cas12a_NT-ssODN and KI_Cas12a_T-ssODN, respectively. The asterisk symbols *,*** and **** indicate statistical significance at *P* = 0.0214, *P*=0.0001 and *P* < 0.0001, respectively (one-way ANOVA, multiple comparison). The control primers used for amplification from both the control and experimental groups, followed by Sanger sequencing results, were monitored for efficiency of programmed CRISPR editing firstly by TIDE followed by NGS and CRISPresso2 of the NGS reads. G,H 5’int-F and 3’int-R are genome-specific primers, and 24nt-6stp-F and 24nt-6stp-R are transgene-specific primers. From the findings, we confirmed that programmed CRISPR/*As*Cas12a gene editing was active in schistosome eggs, and higher than for *Sp*Cas9 by TIDE analysis (ANOVA, ****, *P* 0.0001, n = 12). The estimation of CRISPR efficiency from the 12 biological replicates (each black dot in panels G and H) KI_Cas9 (wide range of CRISPR efficiency from 0.46-2.23 among 12 biological replicates) was higher than for KO based treatments. The estimated CRISPR efficiency in schistosome eggs (LE) from CRISPR/*As*Cas12a (green dots in panels G and H, *n* = 8) was at least 1.5% in most samples. There were no statistically significant differences in CRISPR efficiency among the KI_Cas9 vs. KI_Cas12a (panel H) from Sanger sequencing along with TIDE analysis.

### Quantification of transgene copy number by external standard curve

The external standard curve design that we employed to estimate transgene copy number in genomics DNA of the schistosome eggs (Fig 2E and F). The standard curve on simple linear regression (y = −4.0964ln[10] + 44.821, R^2^ = 0.9923) was established with logarithm-10 transformed initial DNA input, plasmid pCR4, which encodes the six stop codon cassette (Ittiprasert *et al*., 2019), as the dependent and the Ct value from the qPCRs as the independent variable (not shown). We estimated the transgene copy number on the 5’ and 3’ of KI site in *ω1* by converting the measured Ct values to log copy numbers using the equations obtained from the standard curves. Subsequently, dividing the means by the antilog of each result, we arrived at an estimate of the transgene copy number. A 790 bp product amplified for the control wildtype sequence and amplicons of 235 bp and 182 bp for the 5’ and 3’ flanking regions, respectively, of the KI genomic DNA containing the six stop codon transgene were analyzed, which revealed 3’-side transgene integration values of 3.61±0.25, 4.44±0.84 and 4.19±1.22 from the KI_*Sp*Cas9, KI_*As*Cas12a-NT-ssODN and KI_*As*Cas12a-T-ssODN treatment groups, respectively (*P* = 0.4057, = 0.6264 and = 0.9155; NS; one-way ANOVA, multiple comparisons, 95% CI for differences of the means) (Fig 2E). However, the presence of asymmetrical HDR was revealed by significant differences among the copy number values for the 5’ side transgene integration estimates: 2.72±0.42, 5.06±0.54, and 1.64±0.39 for the KI_*Sp*Cas9, KI_*As*Cas12a-NT-ssODN and KI_*As*Cas12a-T-ssODN groups, respectively (Fig 2F; *P* = 0.0214, *P*=0.0001 and *P* ≤ 0.0001, *,***, ****, respectively; one-way ANOVA, multiple comparisons).

### Estimation of CRISPR efficiency of *Sp*Cas9 and *As*Cas12a

RNP complexes of *Sp*Cas9-sgRNA and *As*Cas12a-CRISPR RNA (crRNA) targeting *ω1* were delivered to the LE by electroporation (Ittiprasert *et al*., 2019). The efficiency of programmed mutation was assessed by TIDE analysis of Sanger sequence chromatograms (Gier *et al*., 2020; Jacobsen *et al*, 2020), followed by targeted amplicon NGS and analysis of the reads using CRISPResso2 (Clement *et al*, 2019; Pinello *et al*, 2016). Twelve independent biological replicates were carried out by TIDE analysis. *As*Cas12a programmed KO led to significantly more indels than were induced with *Sp*Cas9 (*P* < 0.001, paired, two-tailed *t* = 7.450, df = 11) (Fig 2G). The mean CRISPR efficiency for KO with *Sp*Cas9 was 0.89%±0.43 (range, 0.30-1.31%) and 1.83%±0.29 (range, 1.39-2.54%) with *As*Cas12a, when compared with the control (mock) groups. Programmed mutation by *Sp*Cas9 was more efficient in the presence of donor ssODN, i.e., KI *versus* KO groups, 1.43±0.4, range 0.51-2.31% (Fig 2G and H). By contrast, differences in mutation efficiency were not apparent between the KO and KI groups using *As*Cas12a; 1.75%±0.63 (range 1.11-2.69%) and 1.54%±0.49 (range 1.82-2.20%) for *As*Cas12a-NT ssODN and *As*Cas12a-T ssODN, respectively. For deeper analysis, both on target KO and KI efficiency also was investigated by CRISPResso2 from pooled samples by target amplicon reads by MiSeq 2×250 bp configuration (Illumina).

### Estimates of indels from NHEJ and/or HDR using NGS reads from targeted amplicons

Pooled gDNAs (above) also were used as templates for targeted amplicons for NGS and CRISPResso2 employed to assess the efficiency of transgene KI from HDR in the NGS reads. Comparison of the mutations among all four copies of the *ω1* gene also were undertaken. From whereas TIDE analysis using Sanger direct sequencing chromatograms was used above to estimate % efficiency of CRISPR manipulations in individual biological replicates, here targeted amplicons from pools of gDNAs from the biological replicates was used to acquire libraries of reads (Amplicon EZ, Genewiz). More than 150,000 reads from each NGS library were obtained. Analysis of the reads using CRISPResso2 provided increased sensitivity of detection of programmed gene editing than TIDE, as expected (Sentmanat *et al*, 2018). Specifically, there were 12.26%, 1.66%, 0.12% modified sequences (NHEJ) for KO-*Sp*Cas9 and 9.27%, 1.33% and 0.28% of NHEJ for KO-*As*Cas12a, when compared with the WormBase Parasite reference sequences (identical amplicon from Smp_334170.1, Smp_334170.2, Smp_334240), Smp_333930 and Smp_333870, respectively (Fig 3A and B). A small number of reads were assessed to be ambiguous by CRISPResso2, 0.49% and 0.93% for KO-*Sp*Cas9 and KO-*As*Cas12a, respectively (Fig 3A and B). Deletions were the only type of indel detected, for both *Sp*Cas9 and *As*Cas12a. In like fashion to other studies (An *et al*, 2020; Dumeau *et al*, 2019; Kissling *et al*, 2019), deletion of large sizes resulted from programmed *As*Cas12a editing (from - 2 to 26 bp deletion size) at the programmed DSB, compared to −3 bp at the DSB programmed from *Sp*Cas9 (Fig 3C, D provide representative allele sequences). Both enzymes showed a preference to mutate the reference Smp_334170 copy, and also Smp_334240, rather than the Smp_333930 and/or Smp_333870 copies of omega-1. The numbers of indels arising from NHEJ with *Sp*Cas9 and with *As*Cas12a both increased in the presence of a ssODN donor repair template. The total NHEJ were 15.67%, 28.71% and 21.43% from KI-*Sp*Cas9-NT-ssODN, KI-*As*Cas12a-NT-ssODN and KI-*As*Cas12a-T-ssODN, respectively (Fig 4A, B and C). Deletion size as determined by CRISPResso2 for the KI-*Sp*Cas9 group was larger than for KO-*Sp*Cas9, increasing up to 13 bp length deletions. Similarly, base deletion size from KI-*As*Cas12a was larger than KO-*As*Cas12a with deletion of 42 bp detected (Fig 4D, E and F). Levels of ambiguous results were estimated at 1.66%, 1.03% and 0.73% from KI-*Sp*Cas9, KI-*As*Cas12a-NT-ssODN and KI-*As*Cas12a-T-ssODN, respectively.

**Figure 3.**
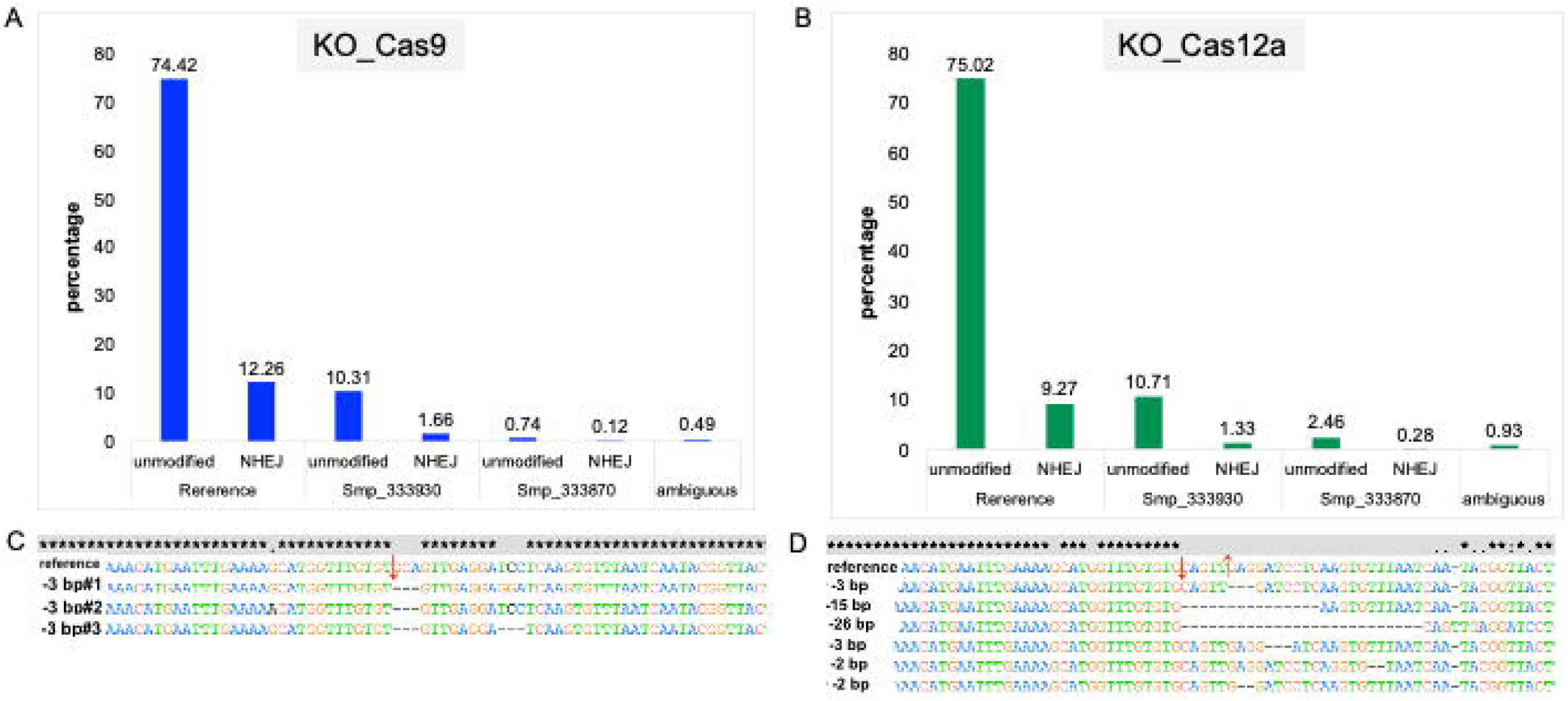
NHEJ assessment from *S. mansoni* eggs at the target *ω1* gene following delivery of *Sp*Cas9 and *As*Cas12a RNP complexes by electroporation. CRISPResso2 analysis of the Illumina sequence reads revealed non-homologous end joining (NHEJ) mutations at targeted loci in schistosome eggs. Allele-specific editing outcomes of CRISPR-*Sp*Cas9 and -*As*Cas12a gene editing in schistosome multiple copies of *ω1* Smp_334170.2, Smp_334170.1, Smp_334240 (100% gene sequence identical; Fig S1B). Mutagenesis frequencies for all loci including Smp_333930 and Smp_333870 were included in CRISPResso2 analysis (Fig S1B). The results were from NGS libraries of pooled genomic DNA from seven biological replicates of KO_Cas9 (Panel A and C) and KO_Cas12a (Panel B and D). The sequence reads assigned to reference genes, Smp_333930 and Smp_333870 loci by the CRISPresso2 to achieve accurate quantification of programmed mutation of the multiple copies of *ω1* from *Sp*Cas9 and *As*Cas12a. The bar chart shows the assignment of each read to the wild type (unmodified) and NHEJ with 12.26%, 1.66% and 0.12% (total 14.04%) from KO_Cas9 on references and other two loci, respectively (panel A). Panel B shows the assigned NHEJ with 9.27%, 1.33% and 0.28% (total 10.88%) from KO_Cas12a on references and two highly identical copies of *ω1* (panel B). The majority of NHEJ indels from both nucleases were nucleotide substitutions followed by gene deletions (Panel C and D). Mutations were induced in all copies of *ω1*; however, most editing was seen on the reference target site. Only deletions were revealed in this experiment, 3 nt deletion downstream from expected cut side (red arrow) was the majority of NHEJ mediated from *Sp*Cas9 (Panel C). The larger deletion from 2 to 26 nt at downstream of expected sticky end cut sites (red arrows) were revealed from NHEJ. CRISPResso2 interpreted ≤ 1.0% of the reads as ambiguous. Ambiguous alignments: 0.49% and 0.93% that could not be attributed uniquely to one of these loci are shown in the right-hand side bar of each graph.

**Figure. 4.**
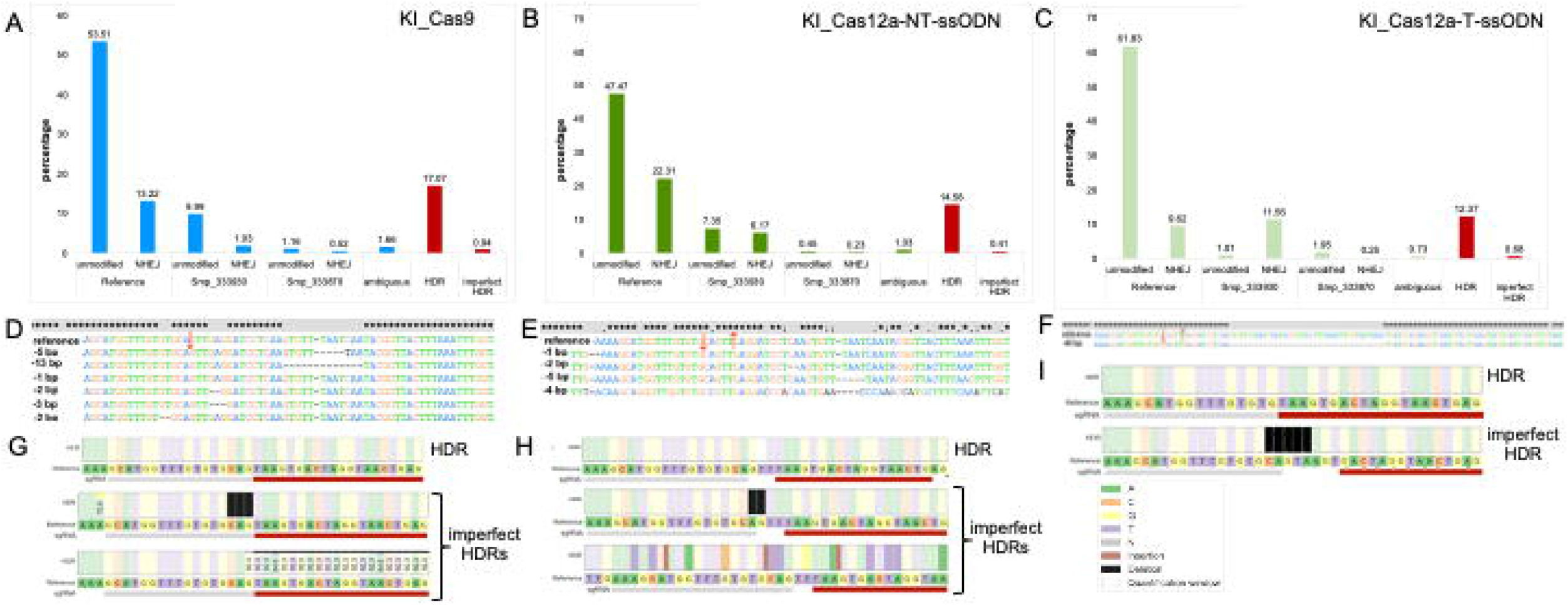
HDR and induction of NHEJ at the *ω1* target in the presence of donor ssODN templates. Panels A to C show the percentage of NHEJ from schistosome eggs that introduced RNP along with ssODN template were higher in the presence of ssODN donor templates than in their absence. There were totals of 15.67% (13.22+1.16+0.52), 28.71% (22.31+6.17+0.23) and 21.43% (9.62+11.56+0.25) editing efficiency in the +KI_Cas9, KI_Cas12a_NT-ssODN and KI-Cas12a_T-ssODN groups, respectively (blue or green bars in each graph). Most mutations were substitutions and deletions located downstream of the programmed double stranded breaks (DSB), with the DSBs indicated by red arrows (Panel D, E and F). A variety of sizes of base deletions from 1 to 15 bp in length were seen in the KI_Cas9, KI_Cas12a_NT-ssODN groups while large deletions, up to 40 bp, were evident as major deletions detected in the KI_Cas12a_T-ssODN group. The percentage of perfect HDR in the three experimental groups, KI_Cas9, KI_Cas12a_NT-ssODN and KI_Cas12a_T-ssODN were 17.07%, 14.58% and 12.37%, respectively (red bars, Panel G, H and I). In addition, imperfect HDRs occurred in all KI treatment groups, likely as the result of interruption of 5’ end HDR gene repair by competing NHEJ activity with the result that deletion of 2-5 nt were seen along with donor transgene insertion (black bars in Panel G, H and I); the frequency of this event was 0.94, 0.41 and 0.68% among the KI_Cas9, KI_Cas12a_NT-ssODN and KI_Cas12a_T-ssODN groups.

To estimate the HDR events, we included the donor sequence as a parameter in the CRISPResso2 analysis. More HDR events mediated by CRISPR/*Sp*Cas9 were detected than with CRISPR/*As*Cas12a; 17.07% *vs* 14.56% and 12.37%, in like fashion to our earlier findings (Santos *et al*, 2021). However, in *Sp*Cas9 sample, more imperfect HDR was seen (0.94%) than in the *As*Cas12a groups (0.41% and 0.68%) (Fig 4A, B, C, F, G and H). Frequently, imperfect HDR event alleles from both Cas nucleases were missing several residues (2-5 nt) on the 5’ side of the programmed DSB. We note that this mutation repair outcome may have impacted quantification of the 5’ KI transgene copy number, especially in KI-*As*Cas12a-T-ssODN samples which contained 5 nt imperfect HDR repair on the six stop codon cassette (primer probing sequence). Imperfect HDR at the 5’ side might result from synthesis-dependent strand annealing (Straume *et al*, 2020).

### Transcription of ω1 interrupted by programmed mutation by *Sp*Cas9 and *As*Cas12a

Primers specific for the four annotated copies of *ω1* were employed to investigate transcription by real time quantitative PCR (Fig S1C). Relative expression levels of *ω1* mRNA in negative controls and experimental KO or KI eggs were compared with wild type. Significantly reduced levels of *ω1* were seen in eggs following the experimental the CRISPR manipulations for gene KO and KI. Transcription of *ω1* was reduced, as follows: 63.76%±5.57, 76.88%±11.33, 71.81%±10.91, 98.69%±0.44 and 80.04%±0.09 in the KO_*Sp*Cas9, KI_*Sp*Cas9, KO_*As*Cas12a, KI_*As*Cas12a-NT-ssODN and *As*Cas12a-T-ssODN groups, respectively. Notably, LE subjected to KI_*As*Cas12a_NT-ssODN retained <2% ω1 transcript levels of the mock control (Fig 5A), and the level in the KI_*As*Cas12a-NT-ssODN group was significantly lower than the KI_*As*Cas12a-T-ssODN group, KI_*Sp*Cas9-ssODN. Significant differences were not apparent between the KO_*Sp*Cas9 and KO_*As*Cas12a groups, generally. There were no statistic differences in levels of transcription among the negative control groups (Fig 5A). Differences among treatment groups were assessed using ANOVA with multiple comparisons (*P* ≤ 0.01 to ≤ 0.0001; n= 4; F = 11.77).

**Figure 5.**
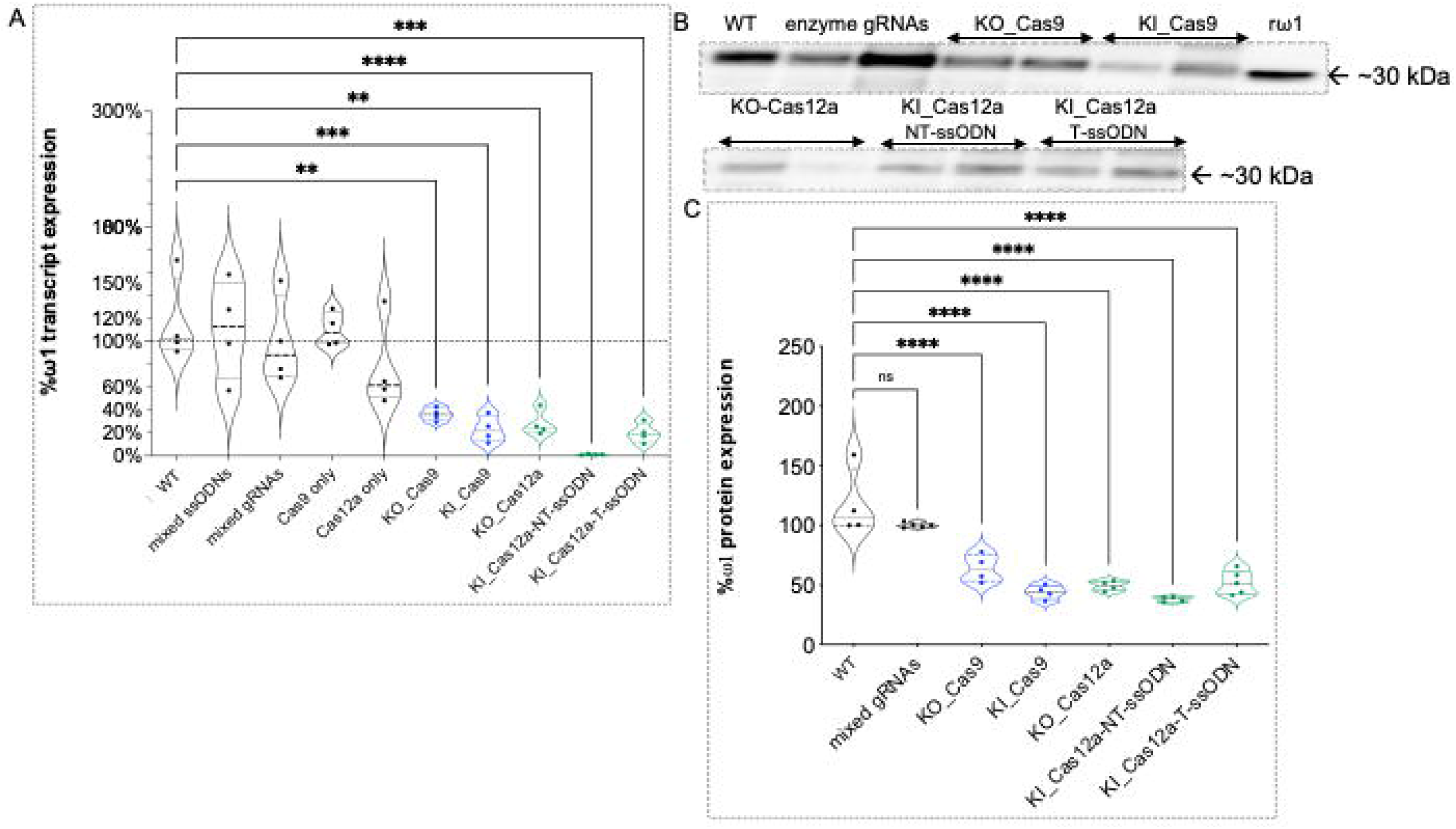
Reduction of the expression of ω1 as assessed by quantitative real time PCR and immunoblot analysis. A Reduction of ω1 transcript levels from KO_Cas9, KI_Cas9, KO_Cas12a, KI_Cas12a_NT-ssODN and KI_Cas12a_T-ssODN eggs compared with control groups after normalization with wild type egg RNA (100% as discontinued line). The means represent four independent biological replicates. B, C Soluble egg antigen; SEA (in duplicates) was sized in SDS-PAGE gels, transferred to PVDF and probed with anti-recombinant omega-1 (rω1) rabbit IgG. Levels of ω1 protein, migrating at ~30 kDa, were reduced in all treatment groups. The percentage of ω1 expression in both KO and KI experimental groups were compared to ω1 levels in wild type (WT) SEA (100%) (panel C) (one-way ANOVA, multiple comparison, *, *P* ≤ 0.05, ***, *P* ≤ 0.001, ****, *P* ≤ 0.0001).

### Protein levels ascertained by western blotting

An anti-rω1 rabbit IgG was used to investigate levels of omega-1 in lysates (soluble egg antigen; SEA) of LE. SEA (200 ng) from the three controls, i.e., the mock, enzyme only, and mixed sgRNAs only groups, and from the KO and KI groups with *Sp*Cas9 or *As*Cas12a was size fractionated by SDS-PAGE and transferred to PVDF and probed with the rabbit antibody (Fig 5B). Following densitometric analysis of the resulting signals, the levels of omega-1 remaining in the SEA were estimated at 64%±11.65, 43.88%±6.02, 49.27%±4.19, 37.9%±2.28 and 52.16%±10 in the KO_*Sp*Cas9, KI_*Sp*Cas9, KO_*As*Cas12a, KI_*As*Cas12a-NT-ssODN and KI_*As*Cas12a-T-ssODN groups, respectively. The differential intensity of ω1expression was not diminished among the sgRNA only groups (Fig 5B and C). By contrast, ω1 protein levels were markedly diminished among all the experimental treatment groups when compared with the controls (*P* ≤ 0.0001) (ANOVA, F = 27.05, *n* = 4). Significant differences among for levels of ω1 among the KO and KI SEA samples were not apparent (Fig 5C).

## Discussion

Given the central role of the egg in disease and in the transmission of schistosomiasis (Costain *et al*, 2018), and given the potential of CRISPR/Cas gene editing to advance understanding of this and other neglected tropical disease pathogens, access to omega-1-negative eggs will provide information to confirm the role of this ribonuclease in disease transmission. Functional knockout (KO) of the *omega-1* gene and the resulting immunologically impaired phenotype have showcased the novel application of CRISPR/*Sp*Cas9 and its utility for functional genomics in schistosomes (Hsu *et al*, 2014; Jinek *et al*., 2012; McVeigh & Maule, 2019; Sankaranarayanan *et al*., 2021). In view of this progress, and the demonstrated tractability of *Sp*Cas9-catalyzed KO of the *omega-1* gene, here we compared the performance of a second RNA-guided Cas nuclease, the *As*Cas12a nuclease of *Acidaminococcus* sp., in terms of efficiency and precision with those for *Sp*Cas9. We now report that *As*Cas12a and *Sp*Cas9 both provide tractable routes for RNA-guided programmed mutation of the genome of the schistosome egg.

We targeted the well-known omega-1 extracellular T2 ribonuclease secreted by the mature egg of *S. mansoni*, which is a key modulator of pathogenesis and transmission (Costain *et al*., 2018; Everts *et al*, 2009; Schwartz & Fallon, 2018; Steinfelder *et al*, 2009). Guide RNAs for both *Sp*Cas9 and *As*Cas12a were designed to anneal to protospacer regions that are essentially at the same target site; the programmed DSB were separated by only three nucleotide residues, on the non-target strand of the target gene. The optimal protospacer adjacent motif for *As*Cas12a, TTTV (Swarts & Jinek, 2019; Zetsche *et al*., 2015), is expected to occur abundantly in the AT-rich genome of *S. mansoni* (Protasio *et al*, 2012). Programmed DSB catalyzed by *As*Cas12a results in a staggered strand break of the target gene sequence, which can enhance efficiency of homology directed repair (HDR) (Swarts & Jinek, 2019). For RNPs assembled from the nuclease and cognate crRNA, *As*Cas12a was significantly more efficient for gene knockout than *Sp*Cas9, as assessed by TIDE, presumably reflecting non-homology end joining (NHEJ)-catalyzed repair and gene silencing.

We also investigated HDR-mediated gene editing with *Sp*Cas9 and *As*Cas12a delivered by electroporation in the presence of the RNPs and donor repair templates. Each donor ssODN template for *Sp*Cas9- and *As*Cas12a-programmed KI included a transgene cassette of 24 nt in length encoding six stop codons (Ittiprasert *et al*., 2019) but in which the homology arms differed slightly on the non-target strand due to the discrete positions of the predicted DSB for each nuclease. For programmed KI catalyzed by CRISPR/*As*Cas12a, we also included as the donor template, the reverse complement sequence of the six stop codon cassette flanked by the target stand sequence as the homology arms. This additional donor was included in our analysis, given concern with respect to the known indiscriminate t*rans*-activity by activated *As*Cas12a against single stranded DNA(Chen *et al*., 2018), and especially for the ssODN with NT homology arms which retains the PAM and partial (~18 nt) protospacer sequence (Swarts, 2019; Swarts & Jinek, 2018, 2019)

RNA-guided *Sp*Cas9 cleavage led to higher levels (~4% higher) of precise knock-out compared to that obtained with *As*Cas12a but markedly less precise knock-in (~6-13%). Deletions were the frequent mutation type seen with both *Sp*Cas9 (deletions of 3 nt in length) and *As*Cas12a (deletions of 2 to 26 nt in length) due to NHEJ repair. Using conditions optimized previously for *Sp*Cas9 KI (Ittiprasert *et al*., 2019), CRISPR/*Sp*Cas9 induced higher (~3%) HDR levels than CRISPR/*As*Cas12a to insert the transgene for both *As*Cas12a-NT-ssODN and *As*Cas12a-T-ssODN donors. Both *Sp*Cas9-and *As*Cas12a-edited eggs exhibited reduced levels of transcription and expression of omega-1. A spectrum of chromosomal alleles was apparent following programmed knock-in of the ssODN repair templates. Both target strand and non-target strand donors were provided. Whereas accurate HDR was catalyzed by both *Sp*Cas9 and *As*Cas12a, precise repair accounted for only a minority of the payload integrations. In some cases, the two sides of the DSB programmed by *As*Cas12a were repaired by different pathways since mutant alleles that included only the left or only the right side of the transgene were observed (Canaj *et al*, 2019). Moreover, accurate HDR was significantly more frequent in the presence of the non-target strand compared with target strand donor.

These findings may have been expected based on reports with other cells and species that indicate that integration of a repair template is not always precise (Paix *et al*, 2017; Ranawakage *et al*, 2020). In addition, *Sp*Cas9 may lead to higher levels of HDR correction than *As*Cas12a in some contexts including in HEK293T and pluripotent stem cells where mis-integrations patterns indicate that two sides of a DSB are repaired by separate cellular repair pathways including the involvement of microhomology as a driver of the mis-integration (Santos *et al*., 2021).

In addition to its guide RNA programmed gene editing and DSB activity, *As*Cas12a and related Type 2 subtypes V and VI nucleases display target substrate bound, non-specific *trans*-activity against ssDNA, dsDNA and/or RNA (Fuchs *et al*, 2019; Nguyen *et al*, 2020). Accordingly, we speculated that activated *As*Cas12a associated with the target exon 1 site in *omega-1* may digest the donor ssODN provided as the repair template. If so, investigation of comparative efficiency of *As*Cas12a *versus Sp*Cas9 catalyzed knock-in would have been compromised. To investigate this possibility, we performed a series of assays *in vitro* with both non-activated and activated *As*Cas12a. Under the conditions and time frame of the reactions - commercially sourced *As*Cas12a, 37°C, 100 mM NaCl, and 120 min reaction duration, we failed to detect degradation of the ssODNs. Although our assays obviously cannot exactly reproduce the physiological environment of the transfected schistosome eggs *in vivo*, we failed to detect indiscriminate *trans*-activity against the ssODNs. However, lack of apparent *As*Cas12a trans-activity *in vitro* does not preclude that indiscriminate *trans*-activity does not take place within the nucleus of the cells within the egg. The RNP may not have been retained at the programmed target site, perhaps degraded by cellular enzymes and/or displaced by the cellular repair machinery following cleavage before *trans*-activity damaged the donor ssODN or other phenomena including protection of cellular DNA by histones. Nonetheless, both *Sp*Cas9 and *As*Cas12a delivered similar knock-in performance in these schistosome eggs.

Whereas enhanced mutation efficacy of *As*Cas12a over *Sp*Cas9 has been reported, the reverse outcome also has been seen (Hur *et al*, 2016; Kim *et al*, 2016). Comparisons among studies are not straightforward because of differences in species, target genes, delivery method, and temperature, among other confounders. In addition, there are three widely used orthologues of *As*Cas12a, *Fn*Cas12a from *Francisella novicida U112, Lb*Cas12a from *Lachnospiraceae bacterium ND2006*, and *As*Cas12a from *Acidaminococcus* sp. *BV3L6* and these have been tested in diverse eukaryotes (Mustafa & Makhawi, 2020). Other orthologues have also been studied (Tóth *et al*, 2018; Zhang *et al*, 2021). *As*Cas12a in active at 37°C whereas other orthologues are catalytically active at lower temperatures, which has enabled studies with zebrafish (Liu *et al*, 2019), insects, and maize (Zhang *et al*., 2021), and environmental and diagnostic biosensing (Chen *et al*., 2018; Ma *et al*, 2020; Nguyen *et al*, 2021).

*As*Cas12a provided enhanced programmed knockout activity compared to *Sp*Cas9 at a specific target site in the *omega-1* gene locus but both exhibited similar performance for HDR based transgene insertion. Although these findings do not enable extrapolation to performance of *As*Cas12a for CRISPR/Cas-based gene editing of other schistosome genes, they demonstrate that Cas12a is active in schistosomes and expand the toolkit for functional genomics in helminth parasites (Sankaranarayanan *et al*., 2021; Zamanian & Andersen, 2016). Optimization of the *As*Cas12a-based approach with regard to ratios of guide RNA:*As*Cas12a and/or ssODN, format of donor(s), and/or temperature will be informative (Chen *et al*., 2018; Hendel *et al*, 2015; Swarts & Jinek, 2018). We anticipate that *As*Cas12a will find increasing use in functional genomes for AT-rich genomes, including schistosomes and other platyhelminths (International Helminth Genomes, 2019). Since transgene insertion into the zygote within the newly released egg has already been described and shown to enable establishment of lines of transgenic schistosomes, advances with gene editing approaches to this key developmental stage of the schistosome life cycle portend progress in the near future with informative gain-of-function lines of transgenic schistosomes.

## Materials and Methods

| Reagent/Resource | Reference or Source | Identifier or Catalog Number |
| --- | --- | --- |
| **Experimental Models** |  |  |
| *Schistosoma mansoni* eggs | Schistosomiasis Resource Center, Biomedical Research Institute, Maryland, USA |  |
| **Recombinant DNA** |  |  |
| recombinant omega-1 protein | Dr. Ron Hokke | Gifted from Dr. Ron Hokke |
| **Antibodies** |  |  |
| Polyclonal antibody against recombinant omega-1 protein | Research Resource Identifier (RRID) | AB_2889886 |
| goat anti-rabbit IgG (H+L) horse radish peroxidase conjugate | Life Technologies, Thermo Fisher Sciences | Cat # 31460 |
| **Oligonucleotides and sequence-based reagents** |  |  |
| PCR primers | This study | Supplement Table S2. They all synthesized from IDT |
| ssODN | Ittiprasert *et al.*, 2019 and this study | Ultramer DNA Oligos, Synthesized from IDT |
| synthetic guide RNA | Ittiprasert *et al.*, 2019 and this study | Synthesized from IDT |
| **Chemicals, enzymes and other reagents** |  |  |
| Alt-R® CRISPR-Sp HiFi Cas9 enzyme | Integrated DNA Technologies | 1081060 |
| Alt-R® CRISPR-AsCas12a enzyme | Integrated DNA Technologies | 1081068 |
| DNAzol® ES reagent | Molecular Research Center | DN127 |
| RNAzol® RT Reagent | Molecular Research Center | RN190 |
| Opti-MEM | Fisher Scientific | Gibco™ 31985062 |
| DMEM | Fisher Scientific | 88364 |
| fetal bovine serum | Fisher Scientific | Gibco™ 26140079 |
| 100× Penicillin-Streptomycin-Fungizone | Fisher Scientific | SV30079.01 |
| NucleoSpin® Gel and PCR clean up | Takara | 740609 |
| SsoAdvanced Universal SYBR Green Supermix | Bio-Rad | 1725271 |
| RIPA buffer | Millipore-Sigma | R0278-500ML |
| Bradford reagent | Bio-Rad | 5000201 |
| SDS-PAGE polyacrylamide gradient gels, 4-12% gel, w/v | Thermo Fisher Scientific | NP0322PK2 |
| Maxima First Strand cDNA Synthesis Kit for RT-qPCR, with dsDNase | Thermo Fisher Scientific | K1672 |
| Peroxidase substrate; Clarity™ Western ECL Substrate | Bio-Rad | 1705062S |
| GoTaq® Green Master Mix | Promega | M712 |
| NEB buffer 2.1 | NEB | B720S |
| Proteinase K | Thermo Fisher Scientific | EO0491 |
| Agilent DNA 1000 Kit | Agilent Technologies | 5067-1505 |
| Agilent small RNA kit | Agilent Technologies | 5067-1548 |

| Software |  |  |
| --- | --- | --- |
| CHOPCHOP | http://chopchop.cbu.uib.no/ |  |
| TIDE | https://tide.deskgen.com/ |  |
| CRISPREsso2 | https://crispresso.pinellolab.partners.org/submission |  |
| Snap Gene | https://www.snapgene.com/ | Academic Subscription Plan |
| GraphPad Prism | Prism software | Monthly Subscription Plan |
| Other |  |  |
| CFX96 Touch Real-Time PCR | Bio-Rad |  |
| ElectroSquarePorator | BTX | ECM830 |
| Chemi-Doc MP System | Bio-Rad |  |
| BioAnalyzer | Agilent Technology |  |

### Ethics statement

Mice experimentally infected with *S. mansoni*, obtained from Schistosomiasis Resource Center (SRC) at the Biomedical Research Institute (BRI), MD were housed at the Animal Research Facility of the George Washington University (GWU), which is accredited by the American Association for Accreditation of Laboratory Animal Care (AAALAC no. 000347) and has an Animal Welfare Assurance on file with the National Institutes of Health, Office of Laboratory Animal Welfare, OLAW assurance number A3205-01. All procedures employed were consistent with the Guide for the Care and Use of Laboratory Animals. The Institutional Animal Care and Use Committee (IACUC) at GWU approved the protocol used for maintenance of mice and recovery of schistosomes. Swiss albino mice were euthanized seven weeks after infection with *S. mansoni*, livers were removed at necropsy, and schistosome eggs recovered from the livers (Dalton *et al*, 1997). The ‘liver eggs, termed LE’ population is expected to include eggs of a spectrum of age ranging from newly released by the female schistosome through to mature eggs containing a fully developed miracidium (Jurberg *et al*, 2009). LE were maintained in DMEM medium supplemented with 20% heat-inactivated fetal bovine serum (FBS), 2% streptomycin/penicillin at 37°C under 5% CO_2_ in air for 18-24 hours before use in gene editing assays (below) (Dalton *et al*., 1997; Ittiprasert *et al*., 2019; Lu *et al*, 2021; Mann *et al*, 2010; Mann *et al*, 2014).

### CRISPR/Cas target design and single stranded DNA donors

The *ω1*-gene specific gRNA for *Sp*Cas9 and *As*Cas12a were designed to target all four copies of the hepatotoxic ribonuclease *omega-1* gene. All four are located on *S. mansoni* genome update annotation of version 7 (updated 16-September-2021 on WormBase ParaSite, https://parasite.wormbase.org/Multi/Search/Results?species=all;idx=;q=Hepatotoxic%20ribonuclease%20omega-1;site=ensemblunit&filter_species=Schistosoma_mansoni_prjea36577); Smp_334170.1, Smp_334170.2, Smp_334240 and Smp_333930 by an online CRISPR design tool, CHOCHOP (Labun *et al*, 2016; Labun *et al*, 2019; Montague *et al*, 2014). There are 99-100% sequence and coding sequence identity among those copies (Fig S1A), which encoding an enzyme of 115amino acids in length, respectively. These *ω1* copies are located on chromosome 1 at nucleotide (nt) positions 3980364 to 3984675, 3992964 to 3995246, and 3908953 to 3911250, respectively. The guide RNA (gRNA), which targets exon 1 of *ω1* complimentary to the catalytic active site histidine [ within FRKHEFEKHGLCAVEDPQV] codons, for the gRNAs programming both *Sp*Cas9 and *As*Cas12a. In addition, the gRNAs exhibited similarly high predicted CRISPR efficiency scores along with an absence of predicted off-target cleavages, as predicted using the CHOPCHOP algorithm using Smp_334170.2 as template (Fig 1A and B). Specifically, the sequence of the gRNA used with *Sp*Cas9 was 5’-GCATGGTTTGTGTGCAGTTG-3’, 20 nt in length and that for *As*Cas12a was 5’-AAAAGCATGGTTTGTGTGCA -3’, 20 nt on exon 1 of Smp_334170.2, Smp_334240 and Smp_333930 or exon 4 of Smp_334170.1 (Fig S1A) The expected cut sites of two nucleases are discrete, being 3 and 2 nt apart from each other on the non-target (NT) and target (T) strands, respectively (Fig 1A).

Single stranded oligonucleotide donors (ssODN) were designed to insert the 24 bp encoding six-stop codons into the programmed cleavage target. The homology arms used in the ssODN donor with *Sp*Cas9 was the non-target strand flanking both the 3’ and 5’ sides of the blunt ended DSB, termed here *Sp*Cas9-ssODN. The homology arms of the two donor ssODNs used with *As*Cas12a (cut at 18 nt and 23 nt on NT and T strands, leading to a sticky ended DSB) were either the target stand or the non-target strand sequences (Fig 1B and C), here termed *As*Cas12a-NT-ssODN and *As*Cas12a-T-ssODN, respectively. The complementary reverse of the 24 nt stop codon cassette was used in the *As*Cas12a-T-ssODN donor, aiming for HDR insertion of the stop codons in the translated forward strand of the gene (Fig 1C).

### CRISPR reagents

Alt-R^®^ CRISPR-*Sp*Cas9 and Alt-R^®^ CRISPR-*As*Cas12a reagents (Alt-R^®^ S.p. HiFi Cas9 nuclease V3, synthetic gRNA (sgRNA) (Labun *et al*.; Labun *et al*.; Nishimasu *et al.*), Alt-R^®^ As *As*Cas12a V3 and crRNA), and ssODN HDR donor templates (Ultramer DNA oligos) and Alt-R^®^ *Sp*Cas9 or *As*Cas12a Electroporation enhancer buffers were from Integrated DNA Technologies (IDT) (Coralville, IA). Figure 1C provides the nucleotide sequences of the gRNAs and donor ssODNs.

### Ribonucleoprotein (RNP) assembly and delivery by electroporation

Stock solutions of the gRNAs for use with *Sp*Cas9 and *As*Cas12a and of the ssODNs were prepared at 1.0 μg/μl in Opti-MEM medium (Sigma, St. Louis, MO). Each gRNA was combined with the respective nuclease at 1:1 ratio to form an RNP complex as described (Ittiprasert *et al*., 2019). The mixture was incubated at room temperature for 15 min, after which 3 μl of Alt-R^®^ *Sp*Cas9 or *As*Cas12a electroporation enhancer was added. The mixture was transferred to a chilled 2 mm electroporation cuvette (BTX), containing ~ 5,000 LE in 100 μl Opti-MEM. LE in the presence of RNPs and the other reagents were subjected to square wave electroporation with a single 125 volts, 1 pulse for 20 ms (ElectroSquarePorator, ECM830, BTX San Diego, CA). In KI assays, 6 μg of ssODN was included in the cuvette before electroporation. Following delivery of the RNPs, LE were maintained for 10 days at 37°C, 5% CO_2_ in air. In addition to experimental gene-edited groups, control groups included LE electroporated in Opti-MEM (mock), mixed-Cas enzymes only, mixed-sgRNAs only, and mixed-ssODNs only. Experimental and control groups were otherwise treated similarly.

### PCR to detect a multiple stop codon containing transgene

Genomic DNAs were extracted from LE using the DNAzol^®^ RT reagent (Molecular Research Center Inc, Cincinnati, OH), and the concentration and purity determined by spectrophotometry (Nanodrop 1000, Thermo Fisher Scientific Inc., Waltham, MA). The HDR target site(s) on either the 5’ or 3’ KI transgene was amplified independently with two pairs of primers (Fig 2A): 1) 5’KI: 5’int-F and 24nt-6stp-R primers and 2) 3’KI: 24nt-6stp-F and 3’int-R using the GoTaq DNA polymerase mix (Promega, Madison, WI) and 200 nM of each primer. The thermal cycling included denaturation at 95°C, 3 min followed by 30 cycles of 94°C, 30s, 56°C, 30s and 72°C, 30s and a final extension for 5 min at 72°C. Thereafter, the amplicons (10 μl) were size separated by agarose gel electrophoresis, stained with ethidium bromide, and imaged using the ChemiDoc Imaging System (Bio-Rad, Hercules, CA). The expected sizes for the 5’KI and 3’KI amplicons were 235 and 182 bp, respectively (Fig 2D). A positive control PCR to amplify ~790 bp flanking the DSB was included.

### Real time qPCR to estimate transgene copy numbers

A SYBR green quantitative PCR-based approach was used to estimate transgene copy numbers using two pairs of primers; 5’KI and 3’KI as detailed (Fig 2A and D) with the SsoAdvanced Universal SYBR Green Supermix (Bio-Rad). Briefly, one ng of genomic DNA (gDNA) was amplified with 10 μl of 2× SsoAdvanced Universal SYBR Green Supermix and 200 nM of each primer, using the CFX Connect Real-Time PCR Detection System (Bio-Rad). PCR conditions included denaturation at 95°C, 3 min followed by 30 cycles of 94°C, 30 s, 56°C, 30 s and 72°C, 30 s and a final extension at 72°C for 5 min, with the signals collected at the annealing step in each cycle. Analysis of transgene copy number in the pooled gDNAs from LE involved comparison of the qPCR signals obtained using a 10-fold serial dilution series of known quantities of the 24 nt six-stop codon transgene in 235 bp (5’ integration PCR fragment) ligated-pCR4-TOPO plasmid DNA (size 4,191 bp) to establish a standard curve (Shirvani *et al*, 2020). The standard curve was based on a linear regression generated using logarithm-10 transformed initial DNA input as the dependent variable and the Ct number of the qPCR signal as the independent variable (Ct values in each dilution were measured in triplicate). Ct numbers were used to fit into the linear regression to derive the estimation of transformed DNA amount for the transgene (Lee *et al*, 2006; Yuan *et al*, 2007). The concentration of the 5’KI-transgene ligated-pCR4-TOPO plasmid was measured (NanoDrop 1000) and the corresponding copy number was calculated as follows:

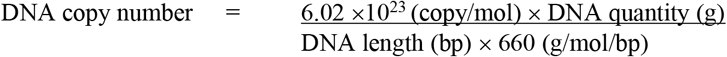

Data were collected from four biological replicates. 5’KI and 3’KI transgene copy numbers from the *Sp*Cas9 and *As*Cas12a treatments were plotted using GraphPad Prism v9 (Fig 2E and F).

### Tracking of INDELs by decomposition (TIDE) analysis

Genomic DNAs from LE exposed to the RNPs were isolated as above. The target *omega-1* locus in the replicates of gDNAs was amplified using a control primer pair (Fig 2A). The amplicon of 790 bp in size from control and experimental groups, both KO and KI focused manipulations, was purified by NucleoSpin^®^ Gel and PCR clean up (Macherey-Nagel, Bethlehem, PA) after which the nucleotide sequence determined by the Sanger method (Genewiz, South Plainland, NJ). Sanger sequence traces from the 12 independent replicates of the experimental groups for both *Sp*Cas9 or *As*Cas12a- (Jacobsen *et al*., 2020) and *Sp*Cas9-catalyzed gene editing manipulations were analyzed and compared with reads from the control (mock) group using the Tracking of Indels by Decomposition (TIDE) algorithm (Brinkman *et al*, 2014; Brinkman & van Steensel, 2019; Meshalkina *et al*, 2020). TIDE provides a rapid and informative assay and also inform decisions on whether proceed to more detailed high throughput, next generation sequencing (NGS) of targeting amplicons (Germini *et al*, 2018).

### On-target amplicon next-generation sequencing

Subsequently, the Amplicon-EZ next generation sequencing approach (Genewiz) was used for deeper coverage of the *ω1* coding sequence mutations and provided > 50,000 reads per sample. Amplicons of 426 bp obtained with the primers bearing Illumina partial adapter sequences at the 5’ end - NGS-F 5’ - TATTGTCAACGGCGTACAGG-3’ and NGS-R 5’-CAATCGGCACTGAGACGCA-3’ - flanking the programmed DSBs for CRISPR/*Sp*Cas9 and CRISPR/*As*Cas12a were sequenced using Illumina chemistry. Partial Illumina adapters were added to 5’ end of the forward and reverse primers forward sequencing read: 5’-ACACTCTTTCCCTACACGACGCTCTTCCGATCT-3’, reverse sequencing read: 5’-GACTGGAGTTCAGACGTGTGCTCTTCCGATCT-3’. The amplicons were purified using the NucleoSpin^®^ Gel and PCR clean up kits from Macherey-Nagel (Bethlehem, PA), after which quantification and qualification of purified band(s) were undertaken using the Agilent DNA 1000 kit and BioAnalyzer (Agilent). Illumina NGS was performed with 2×250 bp sequencing without fragmenting the amplicons. Raw reads in FASTQ format were analyzed for mutations resulting from NHEJ and HDR by CRISPREsso2 (Clement *et al*., 2019) using the reference *ω1* sequences, Smp_334170 and Smp_333930 including Smp_333870 (this latter is annotated as an uncharacterized protein but shares >99% identity) (Fig S1B). The default parameters were used to estimate indel and HDR events. The sequence reads from the Amplicon-EZ runs are available at GenBank Bioproject PRJNA415471, BioSample SAMN07823308, SRA study SRP126685, accessions SRR: 13498338-13498342, 15971309-15971320, and 6374209-6374210.

### Quantitative real time PCR

Total RNA of LE was isolated using the RNAzol^®^ RT reagent (Molecular Research Center) according to the manufacturer’s instruction. RNA concentration and purity were determined (Nanodrop 1000 spectrophotometer, Thermo Fisher). Ten nanograms of RNA was treated with DNase I to remove residual DNA then converted to cDNA by Maxima First Strand cDNA Synthesis Kit (Thermo Fisher Scientific). The ω1-specific product was amplified using SsoAdvanced Universal SYBR Green Supermix (Bio-Rad) on CFX96 real time PCR (Bio-Rad) using the primers as described (Ittiprasert *et al*., 2019), ω1-RT-F: 5’-GGTTACTTTAAATTTGGTA-3’ and ω1-RT-R; 5’-AGTTTCCAAGGAACGGGCAG-3’. These primers amplify all annotated ω1 transcript copies; Smp_334170.1, Smp_334170.2, Smp_334240 and Smp_333930 (Fig S1C). The PCR reaction was denatured at 95°C for 30 s, 40 amplification cycles each consisting of denaturation at 95°C for 15 s and annealing/extension at 60°C for 30 s. The output was analyzed in CFX manager software (Bio-Rad). Relative expression of ω1 was calculated using the 2^−ΔΔCt^ method and normalized to schistosome GAPDH (Smp_0569701) expression using specific primers; forward 5’-ATGGGACATTTCCAGGCGAG-3’, reverse 5’-CCAACAACGAACATGGGTGC-3’ (Livak & Schmittgen, 2001). Data four biological replicates were compared to wild type LE and fold change reported as mean ± SE (*n* = 4).

### Western blot analysis

Soluble egg antigen (SEA) of LE was extracted by mechanical lysis (Funakoshi motorized pestle; Diagnocine, Hackensack, NJ) in RIPA buffer (Millipore Sigma) supplemented with protease inhibitors (Protease Inhibitor Cocktail I, Millipore Sigma), as described (Ittiprasert *et al*., 2019). The lysate was clarified by centrifugation at 13,000 rpm, 15 min, 4°C after which protein concentration was determined by the Bradford method (Kielkopf *et al*, 2020). Ten micrograms of SEA were size-separated through SDS-PAGE polyacrylamide gradient gels, 4-12% gel, w/v (Bolt Bis-Tris Plus, Invitrogen, Thermo Fisher Scientific) before transfer to PVDP membranes (Transblot Turbo Transfer System, Bio-Rad). Polyclonal anti-ω1 IgG antibodies were purified from the rabbit anti-ω1 sera, raised as follows. Two NZ white rabbits, E12983 and E12984, were immunized with 30 μg of recombinant *omega-1* RNase T2 of *Schistosoma mansoni* (rω1) (produced *in planta* by agroinfiltration of leaves of *Nicotiana benthamiana* (Wilbers *et al*, 2017)) in Complete Freund’s Adjuvant and boosted four times at two-week intervals using 15 μg rω1 in Incomplete Freund’s Adjuvant each time. Blood was collected 66 days after the first immunization, IgGs was isolated from sera using an immobilized protein A ligand. The concentrations of the purified antibodies were 1.31 mg/ml (E12983) and 1.51 mg/ml (E12984). These antibodies have been assigned Research Resource Identifier (RRID) AB_2889886 (Bandrowski *et al*, 2016). PVDF membranes were probed with the E12983 IgG antibody, diluted 1:1,000, for 18 h at 4°C in PBS-Tween (0.5%), 5% non-fat powdered milk, washed 3 times in PBS-Tween (wash buffer), and probed with goat anti-rabbit IgG (H+L) horse radish peroxidase conjugate (Life Technologies, Thermo Fisher Sciences), diluted 1:5,000, for 60 min at 23°C with gentle shaking. Following washing (3 times in wash buffer), the membrane was exposed to a peroxidase substrate (Clarity Western ECL Substrate, Bio-Rad) for 5 min at 23°C, after which chemiluminescence was quantified and imaged (Chemi-Doc MP System, Bio-Rad). The Image Lab software (Bio-Rad) was used to quantify the ω1 signals. The percentage of ω1 expression from the experimental and control groups was established by comparison with that of wild type SEA, assigned as 100%.

### Single stranded DNA donors as a potential substrate for indiscriminate *trans* activity of activated *As*Cas12a

Previous reports reveal the *trans*-activity of wild type *As*Cas12a prefers ssDNAs over dsDNAs, and was inactive toward dsDNA in the presence of magnesium (II) ions (Li *et al*, 2020). The *As*Cas12a exhibited not only endonuclease activity on both dsDNA, *cis*- and *trans*-activity (Li *et al*, 2018b). Accordingly, to investigate whether activated *As*Cas12a could degrade the ssODN donors used in this study by indiscriminate *trans*-activity, several *in vitro As*Cas12a-RNP digestion assays were carried out. These included the dsDNAs containing the *As*Cas12a target sequence used in this study and 5’PAM (TTTG) as positive enzymatic activity. A 426 bp dsDNA template was amplified using the NGS study (above) primers. Activity of *As*Cas12a-RNP against the dsDNA template released fragments of 265 and 161 bp (Fig 2B), at 37°C for ≥30 min as predicted in 1 × NEB Buffer 2.1 (500 mM NaCl, 20 mM sodium acetate, 0.1 mM EDTA, 0.1 mM tris(2-carboxyethyl) phosphine (TCEP), 50% glycerol). *As*Cas12a activity was inactivated by the addition of Proteinase K (1 μl at 20 mg/ml) at 56°C for 10 min. Electropherograms of the digestion products were obtained using the BioAnalyzer (Agilent DNA1000) to visualize the size of templates and fragments.

To investigate indiscriminate *trans*-cleavage of ssODNs, similar *in vitro* digestion reaction was carried out, as above, but with the addition of 50 ng of either *As*Cas12a-NT-ssODN or *As*Cas12a-T-ssODN. A mixture of *As*Cas12a-RNP complex in 1 × NEB Buffer 2.1 and ssODN was incubated at 37°C for 30, 60 and 120 min before terminating the reaction by addition of proteinase K at 56°C for 10 min, and visualized as above. The ssDNA ladder (SimPlex Sciences, NJ) ranging from 10-200 bp was included as the single stranded DNA marker. *Trans*-activity of Alt-R *As*Cas12a V3-RNP complex was not apparent against the ssODNs under these conditions *in vitro* (Fig. 2C).

### Statistical analysis

Statistical analysis was performed using GraphPad Prism version 9.1 (GraphPad Prism, La Jolla, CA). Graphs were plotted as the mean with bars to show the standard error of the mean. The differences among groups were assessed by one-way ANOVA. *P* values of ≤ 0.05 were the statistically significant.

### GenBank/EMBL/DDBJ

#### Database accessions

Nucleotide sequences reads of the amplicons are available at Bioproject PRJNA415471, https://www.ncbi.nlm.nih.gov/bioproject/PRJNA415471 and GenBank accession PRJNA415471, BioSample, SAMN07823308, SRA, SRP126685.

## Acknowledgements

We thank Professor Cornelis Hokke, Leiden University Medical Center, The Netherlands for the gift of recombinant *Schistosoma mansoni* omega-1. We thank the Thailand Research Fund, the Royal Golden Jubilee Ph.D. program, and Dr. Dalina Tanyong, Mahidol University, Thailand, for support for CC to perform the work at GWU. This research was funded in whole, or in part, by the Wellcome Trust (award number 107475/Z/15/Z, titled ‘Functional Genomics Flatworms Initiative’, K. H. Hoffmann, principal investigator). For the purpose of Open Access, the authors have applied a CC BY public copyright license to any Author Accepted Manuscript version arising from this submission. Schistosome-infected mice and snails were provided by the NIAID Schistosomiasis Resource Center of the Biomedical Research Institute, Rockville, Maryland through NIH-NIAID Contract HHSN272201000005I for distribution through BEI Resources.

## Author contributions

WI, CC, and PJB designed the experiments, analyzed the data, and drafted the manuscript with input from all the co-authors; CC and WI performed the gene editing focused experiments; VHM, WL, AM, TO, YNA and MMK contributed the parasite materials to the experiments, to data collection, and reviewed and edited the manuscript; PJB and WI supervised the project. PJB, WI and MMK arranged the funding.

## Conflict of interest

The authors declare that do not have competing interests.

## Supporting information

**Figure S1**. **Gene structure encoding the hepatotoxic ribonuclease, omega 1 (*ω1*). At least five copies of the gene are located on chromosome 1, as apparent in the draft sequence of *S. mansoni*, version 7, scaffold SM_V7_1:3,885,722-4,002,365**. The positions of the several *ω1* genes of the schistosome genome (WormBase Parasite, September 2021), including two loci of Smp_334170, Smp_334240, Smp_334070, Smp_3332220, Smp_333930, Smp_333870 and Smp_179960 are shown to for their physical relationship of the copies on the chromosome.

A Gene structures and sizes of Smp_334170.1, Smp_334170.2 and Smp_334240 and the CRISPR/Cas programmed target sites for *Sp*Cas9 (blue arrow) and *As*Cas12a (green).

B All gene copies of *omega-1* share >99% DNA sequence identity; DNA sequence alignment of the *omega-1* orthologues and target amplicon primer indicated as NGS-F and NGS-R. Four copies of the gene, Smp_334240, Smp_334170 and Smp_333930, are reported in Uniprot.

C Alignment of mRNA sequences and primers used for gene transcript analysis (purple).

**Table S1.** The CRISPREsso2 analysis findings from four negative controls: mixed sgRNAs only, *Sp*Cas9 only, *As*Cas12a only, and mixed donors only. There were over 180,000 reads from each library. There was < 1% of background modified reads from all negative control samples.

**Table S2.** Nucleotide sequences of oligonucleotide primers used in the investigation.

## Notes

### Competing Interest Statement

The authors have declared no competing interest.

### Summary of Updates

The modified abstract and title including few edits in the manuscript text.

